# Stromal resistance to castration-induced prostate regression in a mouse model of benign prostatic hyperplasia

**DOI:** 10.1101/2022.12.07.519520

**Authors:** Renyuan Zhang, Shalini Singh, Chunliu Pan, Bo Xu, Jon Kindblom, Shu-Yuan Yeh, Chawnshang Chang, Kevin H. Eng, John J. Krolewski, Kent L. Nastiuk

**Author notes:** these authors contributed equally to this work (Renyuan Zhang, Shalini Singh). Dept of Biology & Interdisciplinary Unit, Data Science & Analytics, Buffalo State College, SUNY. **Address correspondence to:** Kent L. Nastiuk, PhD, Roswell Park Comprehensive Cancer Center, Carlton & Elm Streets, Buffalo, NY 14263, USA. **Email addresses of all authors:**.

## Abstract

Benign prostatic hyperplasia (BPH) is a non-neoplastic proliferative disease producing lower urinary tract symptoms related to the enlarged prostate. BPH is pathologically characterized by hyperplastic growth in both epithelial and stromal compartments. Androgen signaling is essential for prostate function and androgen blockade is the second-line medical therapy to relieve symptoms of BPH. Here we examined the prostates of probasin promoter-driven prolactin (Pb-PRL) transgenic mice, a robust model of BPH that spontaneously develops prostate enlargement, to investigate prostate regression in response to surgical castration. Serial ultrasound imaging demonstrated very uniform self-limited growth of Pb-PRL prostate volume that is consistent with the benign, limited cellular proliferation characteristic of BPH and that contrasts with the highly variable, exponential growth of murine prostate cancer models. Castration elicited only a partial reduction in prostate volume, relative to castration-induced regression of the normal prostate gland. The anti-androgen finasteride induced a diminished reduction of Pb-PRL prostate volume versus castration alone. The limited extent of Pb-PRL mouse prostate volume regression correlated with the initial volume of the stromal compartment, suggesting a differential sensitivity to androgen withdrawal of the epithelial and stroma compartments. Indeed, two-dimensional morphometric analyses revealed a distinctly reduced rate of regression for the stromal compartment in Pb-PRL mice. The myofibroblast component of the Pb-PRL prostate stroma appeared normal, but contained more fibroblasts and extracellular collagen deposition. Like normal prostate, the rate of regression of the Pb-PRL prostate was partially dependent on TGFß and TNF signaling, but unlike the normal prostate, the extent of castration-induced regression was not affected by TGFß or TNF blockade. Our studies show that androgen deprivation can effectively reduce the overall volume of hyperplastic prostate, but the stromal compartment is relatively resistant, suggesting additional therapies might be required to offer an effective treatment for the clinical manifestations of BPH.

## Introduction

Benign prostatic hyperplasia (BPH) causes prostate enlargement due to an increased number of both stromal and epithelial cells. BPH increases with age and is one of the most common urologic diseases among older men [1]. BPH can compress the urethra and result in bladder outflow obstruction [2]. Lower urinary tract symptoms (LUTS), consequently, are common in men with BPH. A variety of surgical and medical therapies relieve LUTS [2, 3]. Conventional alpha-adrenergic receptor blockers are used to relax smooth muscle and thereby relieve obstruction to urinary flow, but they do not shrink the size of the prostate [4]. White and Cabot first described BPH control by castration in the 1890s [5, 6]. Huggins and Stevens reported in 1940 that reduction in prostatic size after castration was due to epithelial cell atrophy in patients with BPH [7]. Medical therapy for LUTS due to BPH includes inhibiting androgen signaling to improve function and prevent disease progression [8]. In order to minimize the many morbid side-effects of complete androgen blockade via castration [9–11], 5-alpha-reductase inhibitors (5ARIs) such as finasteride, have been developed to reduce androgen signaling by blocking conversion of testosterone (T) into tissue-active dihydrotestosterone (DHT) [12]. DHT is a considerably more potent agonist of the androgen receptor (AR) than T, due to both a longer half-life and a higher affinity for AR [13]. Therefore, using 5ARIs to reduce intraprostatic DHT partially blocks androgen signaling, thereby relieving LUTS while minimizing morbid side-effects due to low circulating testosterone [14].

Huggins and Hodges later described the essential role of androgens in prostate neoplastic growth by demonstrating castration controls prostate cancer [15]. Androgen deprivation therapy (ADT) triggers cell death by apoptosis in both normal and hyperplastic prostate and in prostate cancer [16, 17]. While medical management of BPH no longer includes surgical or medical castration, ADT continues to be standard-of-care therapy for patients with advanced prostate cancer, and in these men also relieves co-incident LUTS [18]. In normal prostate, TNF and TGFβ1 signaling are required for castration-induced regression [11, 17]. TNF and TGFβ1 also mediate regression after ADT in prostate cancer mouse models [19, 20]. The potential role of ADT in the treatment of BPH has been complicated by the observations that increased prostate volume does not always correlate with the clinical syndromes of LUTS [2]. While it is likely that novel therapies that effectively reduce prostate volume will reduce symptoms, human trials are required to ultimately determine the clinical effectiveness of such novel therapies.

In this report, we determined the effect of castration on prostate regression in a robust model of BPH, the Pb-PRL mouse. This model develops urinary retention and flow symptoms resembling LUTS [21]. To determine the effects of androgen withdrawal, we have analyzed castration-induced changes in the volumes of the epithelial and stromal compartments using a combination of ultrasound imaging and morphometric techniques. We report that (i) as opposed to the heterogeneity in prostate cancer, the growth of Pb-PRL hyperplastic prostate is uniform and self-limiting; and (ii) similar to normal and neoplastic prostate, castration induces prostate regression in this BPH model. However, our morphometric analyses showed that while the prostate epithelial compartment in this BPH model regressed similar to normal prostate and prostate cancer, the prostate stromal compartment regressed more slowly, resulting in greater prostate volume after castration/ADT.

## Materials and methods

### Animals

All animal studies were approved by the Roswell Park Comprehensive Cancer Center Animal Care and Use Committee (IACUC 1308M). Probasin-driven prolactin (Pb-PRL) transgenic mice were generated as previously described [22] and backcrossed into an FVB background [21]. Non-transgenic littermates served as controls. A prostate-specific conditional Pten-deficient model of prostate cancer (PbCre4 ×Pten^flox/flox^ mice) was created by crossing Pten^flox/flox^ mice with Pb-Cre4 transgenic males [23]. Colonies were housed in environmentally controlled conditions on a 12-hour light/dark cycle with food and water ad libitum. Prostate volumes were determined by imaging, typically monthly, but up to weekly, during development, and up to twice weekly during treatment.

### High-frequency ultrasound imaging

We previously determined three-dimensional (3D) volume reconstruction from high-frequency ultrasound (HFUS) images accurately monitors normal, neoplastic, and hyperplastic mouse prostate volume changes [24]. Briefly, mice were anaesthetized, shaved, and depilated. Anesthesia was maintained with a continuous flow of 1% isoflurane in oxygen mixture while mouse abdomens were imaged. B-mode HFUS images were obtained using the Vevo 2100 micro-ultrasound imaging system (Vevo LAZR; VisualSonics Inc., Toronto) with a 21-MHz linear-array transducer system (LZ250; 13-24 MHz maximal broadband frequency; 21 MHz center frequency, 75 μm axial resolution, 80 μm lateral resolution, 23 mm maximal lateral field of view) in the Roswell Park Translational Imaging shared resource. Images were imported into Amira software (Visualization Sciences Group, Burlington MA), and manually segmented (identifying anatomic structures) followed by 3D mouse prostate volume reconstruction as previously described [24].

### Animal surgery and treatment

Mice were anaesthetized with a continuous flow of 2% to 3% isoflurane in oxygen. Mature (six to nine month old) mice were pre-screened by HFUS imaging to obtain a baseline prostate volume for enrollment. For studies using surgical castration, 20 Pb-PRL animals with a VP volume of about 120 mm^3^ were randomized based on pre-treatment volume into five groups, sacrificing four mice at each time point: 0, 4, 7, 10 and 17 days after castration. Twenty age-matched non-transgenic littermate animals were similarly randomized into five groups as above. All Pb-PRL and littermates were surgically castrated by removal of both testicles as described previously [17]. Mice were then imaged twice weekly. To assess the effect of 5ARIs, mice were injected intraperitoneally (IP) with finasteride (S1197, Selleck chemicals, 25 mg/kg) daily, imaged twice weekly, and sacrificed after 17 days. Cytokine signaling mediating castration was assessed using either a combination of a TGFβR1/ALK5 inhibitor (SB431542, Stemgent, 10 mg/kg) and a p38 inhibitor (BIRB 796, Biotang, 3.33 mg/kg), by daily subcutaneous injection one day prior to castration and every day thereafter for the duration of the experiment. TNF signaling was blocked using etanercept (Enbrel, Amgen, 4 mg/kg) injected IP 3 days and 1 day before castration, and then twice per week after castration. No animals showed signs of distress, including weight loss, during any of the treatments. Ultrasound imaging was performed before castration and twice weekly after castration. Mice were sacrificed 17 days after castration.

### Histological and immunohistochemical analysis

IHC staining was performed on 5-μm sections of formalin-fixed and paraffin-embedded tissue samples. Antibodies against the proliferation marker Ki-67 (Abcam, ab15580; 1:200 dilution) and alpha-SMA (alpha smooth muscle actin, Abcam, Ab21027; 1:200 dilution) were detected using standard or ChromoPlex dual detection reagents on a Leica Bond automated IHC staining platform in the Roswell Park Pathology Network shared resource. For Ki67, images were captured from each stained slide using Olympus BX45 microscope and cellSens software (Olympus, Shinjuku-ku, Tokyo, Japan). The number of Ki-67-stained cells were counted in the epithelial layers of hyperplastic glands from 5 high powered (200X) fields. Three high power fields of the prostate were analyzed from each animal for myofibroblasts (alpha-SMA IHC). Tissues were stained using Masson’s trichrome (Poly Scientific R&D Corporation, cat. no. K037) to visualize collagen. Masson’s trichrome stain was quantified in three high power fields of the prostate from each animal. Collagen content was analyzed using ImageJ software with the color deconvolution plug-in, which separates a three-channel image into three colors to quantitate blue (collagen) staining, as described [25]. The ‘trichrome score’ is determined by multiplying the area of blue-staining by the staining intensity.

### Morphometric analysis

Quantitation of the epithelial and stromal compartment volumes were derived from two-dimensional measurements of tissue areas by manual tracing of epithelium or stroma on digital images of H&E stained tissue sections, using calibrated image J analysis software. Anti-α-SMA IHC staining of myofibroblasts (which form bands surrounding the epithelial glands) was used to guide the identification of ten high-power fields of H&E stained images for each animal. The two-dimensional (2D) areas were normalized to the three-dimensional (3D) volume of the ventral prostate gland using the corresponding HFUS volume measurements, and the following equation:

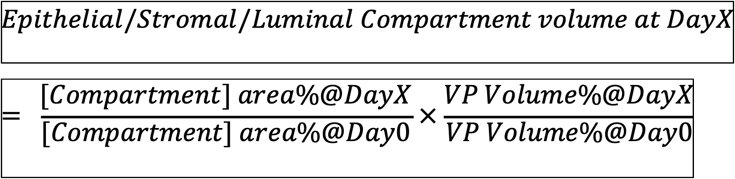

### Statistics

Data are presented as the mean ± SEM. Statistical analyses were conducted using JMP-Pro12 (SAS, Cary, NC, USA). Differences between two means was assessed by Student’s unpaired *t*-test. The compartment specific regression slopes, and accompanying p-values, were determined using a least-squares linear regression model Differences in regression slopes were calculated using a non-linear regression model.

## Results

### Prostate hypertrophy development in Pb-PRL mice

The mass of both the dorsolateral prostate (DLP) and ventral prostate (VP) lobes of transgenic Pb-PRL mice has been reported to increase six-fold versus wild-type prostate lobes during their first year of development [22]. We determined total VP volume of a large cohort of Pb-PRL model mice during development (aging) by serial HFUS imaging. Three dimensional reconstructions of the VP lobe (green) and bladder (yellow) are shown for four representative Pb-PRL mice over 14 weeks in Figure 1A. Determination of VP growth in 44 Pb-PRL mice revealed a remarkably uniform pattern of prostate enlargement; growth kinetics resembled a sigmoidal curve (Fig. 1B), similar to the kinetics of growth in human BPH [1]. VP volume gradually increased to around 20 mm^3^ during the first 15 weeks of life, in accord with the previously reported wet-weight of VP from mice sacrificed at 10 weeks [22]. These VP volumes are comparable to the 21.2 +/-0.4 mm^3^ volume of 6-9 month old non-transgenic FVB littermate (n = 16, data not shown). VP volume then increased rapidly from 20 weeks to 30 weeks of age to ~120 mm^3^ and then growth slowed and plateaued at ~180 mm^3^. In comparison, we previously reported a murine model of prostate cancer (PbCre4 ×Pten^flox/flox^; PTEN-deficient) tumor grew from ~200 mm^3^ at 15 weeks to ~1000 mm^3^ at 30 weeks [20]. In a larger cohort of this prostate cancer model, serial imaging revealed an exponential increase in tumor volume as well as significant animal to animal variability (Fig. 1C). The difference in growth rates for both mouse models is consistent with a fundamental difference between the hyperplastic and neoplastic processes.

**Figure 1.**
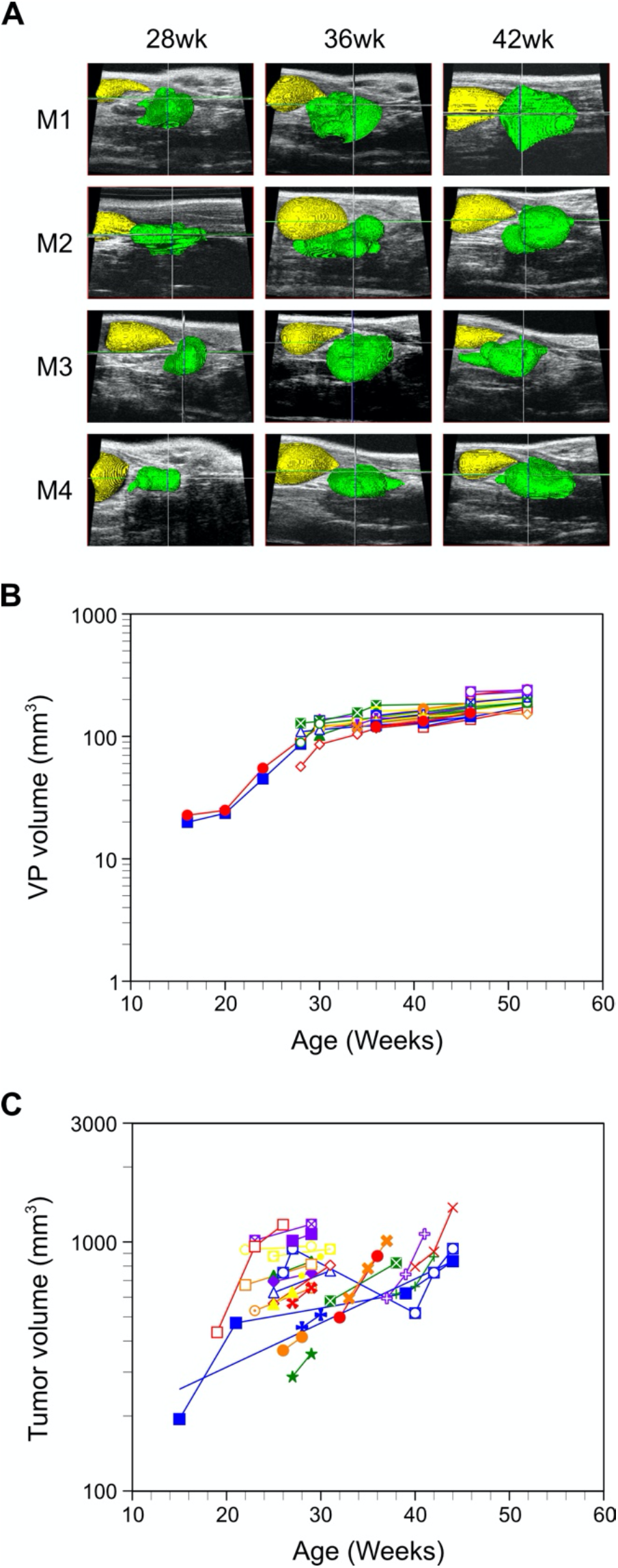
Growth of the ventral prostate lobe in Pb-PRL transgenic mice: sigmoidal growth curve kinetics and limited inter-animal variability. Prostate volumes of Pb-PRL and PbCre4 x PTEN^f/f^ mice were monitored during aging by serial HFUS imaging. **A.** Illustration of 3D-reconstruction of ventral prostate for four Pb-PRL mice acquired 28, 36 and 42 weeks after birth. Visualization of the bladder (yellow) and the ventral prostate (green) was performed by pseudo-coloring. **B.** Kinetics of ventral prostate growth in Pb-PRL transgenic mice. Volumes from the 3D reconstructions are plotted for each animal at the indicated age. Each color represents an individual mouse (N = 44). **C.** Kinetics of prostate tumor growth in prostate-specific PTEN-deficient mice. Volumes from the 3D reconstructions are plotted for each animal at the indicated age. Each color represents an individual mouse (N = 26).

### Castration induced regression in the Pb-PRL model

Next, to determine the kinetics of androgen withdrawal response in this BPH model, we castrated Pb-PRL mice and their non-transgenic (non-Tg) littermates. We then serially determined the VP volume using HFUS imaging (Fig. 2). Previously, we reported that in the normal prostate, castration induces more regression in the VP than the DLP, as the VP contains a higher proportion of epithelial cells than the other lobes [26, 27]. The kinetics of regression were distinct between the non-Tg and Pb-PRL mice. The volume of the non-Tg mice declined in a nearly monotonic fashion until the residual volume approached 10% of the starting volume (Fig. 2C, red line). In contrast, Pb-PRL VP regression occurred in two phases. First, volume decreased more slowly than the non-Tg mice for the first 10 days following castration, to ~60% of starting volume, and then in the second phase, the volume declined even more slowly, to ~50% of the original volume over the next 7 days (Fig. 2C, blue line). Thus, in the BPH model, castration-induced prostate regression is slow, and there remains a significant residual volume.

**Figure 2.**
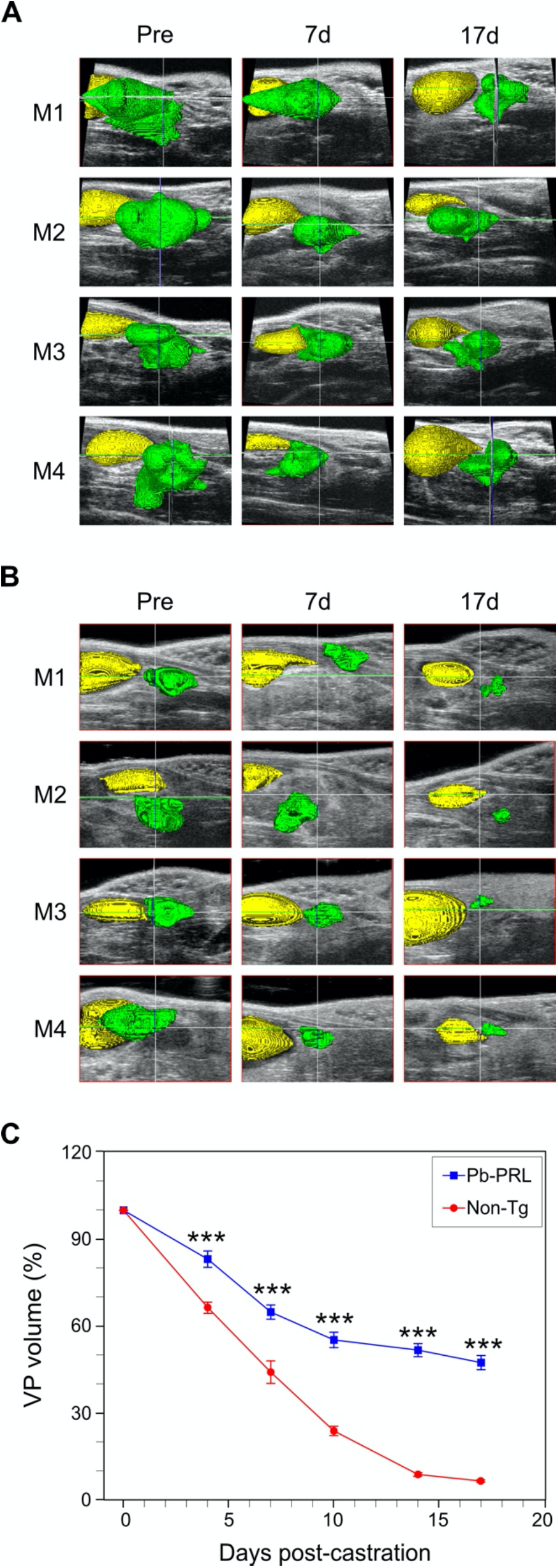
Castration-induced ventral prostate regression reduced in BPH model mice versus normal mice. Pb-PRL transgenic and non-transgenic (Non-Tg) littermate mice were surgically castrated and monitored by serial HFUS imaging. Ultrasound images corresponding to pre-castration (Pre) as well as 7 and 17 days following castration. **A.** 3Dreconstructions of Pb-PRL mice. Visualization of the bladder (yellow) and the ventral prostate (green) was performed by pseudo-coloring. **B.** Similar to **A**, for non-transgenic littermate mice. **C.** Ventral prostate regression following castration in Pb-PRL transgenic mice (blue) and non-transgenic littermate mice (red). Volumes of the 3D reconstructions are plotted at the indicated times following castration. Data points represent mean volume relative to starting volume, +/-SEM (n=4 (non-Tg), n=12 (Pb-PRL). \*\*\**P* <.001; vs non-Tg. Non-Tg and Pb-PRL mice were compared using two-way ANOVA with Tukey’s HSD test.

### Finasteride-induced regression is reduced compared to castration

Finasteride is a 5-alpha reductase inhibitor used to treat LUTS in men with BPH. Finasteride inhibits the of T to the more potent DHT [13]. Since DHT is a more potent androgen, the net effect of finasteride treatment is a reduction in androgen activity in both humans and mice. Therefore, we compared the effect of daily administration of finasteride on the Pb-PRL mouse prostate volume with the effects seen for ADT by castration. We observed that finasteride was less effective in inducing regression of the hyperplastic prostate (Fig. 3).

**Figure 3.**
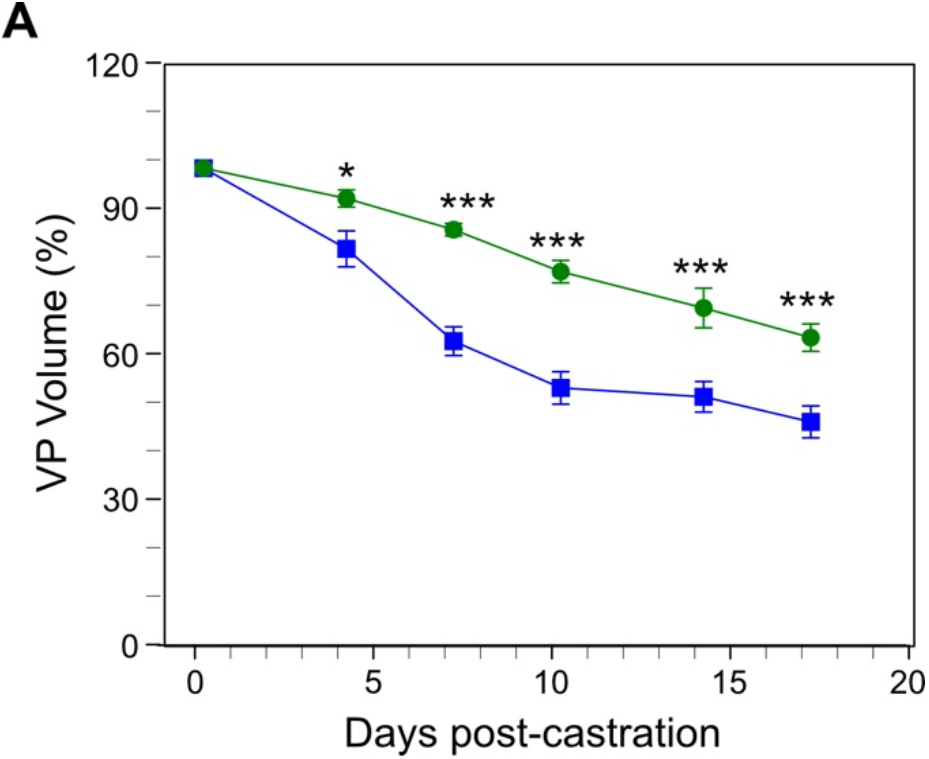
Comparison of ventral prostate regression induced by finasteride versus castration in the Pb-PRL mouse. Pb-PRL mice were surgically castrated (blue line, data from Fig. 2), or left intact and treated daily with finasteride (green line). The ventral prostate of each mouse was monitored by serial HFUS imaging. Segmentation and 3D Amira-reconstruction was performed on the resulting images to determine the volume of the ventral prostate. Data points represent mean volume relative to starting volume, +/-SEM (n=4). *P<.05; \*\*\**P* <.001. Finasteride treated Pb-PRL mice and castrated Pb-PRL mice were compared using two-way ANOVA with Tukey’s HSD test.

### Differences in regression of the epithelial and stromal compartments in Pb-PRL mice

To better understand the cause of the discrepancy in the rates of regression and of the residual prostate volume between the BPH model and normal mice, we characterized the epithelial and stromal prostate compartments of the Pb-PRL mice and of the non-Tg littermates. First, we used Ki67 immunohistochemical (IHC) staining to compare rates of proliferation in the prostate epithelial compartments (Fig. 4A). In both the Pb-PRL and the non-Tg littermate mouse prostate, Ki67 staining was predominantly in epithelial cells. A trend to increased proliferation was observed in the Pb-PRL mice (Fig. 4B), consistent with the hyperplastic epithelial phenotype expected in a model of BPH. However, note that the epithelial glands in the Pb-PRL mouse prostates have relatively little of the undulation and infolding of the glandular epithelial that is characteristic of epithelial proliferation in the hyperplastic glands of human BPH.

**Figure 4.**
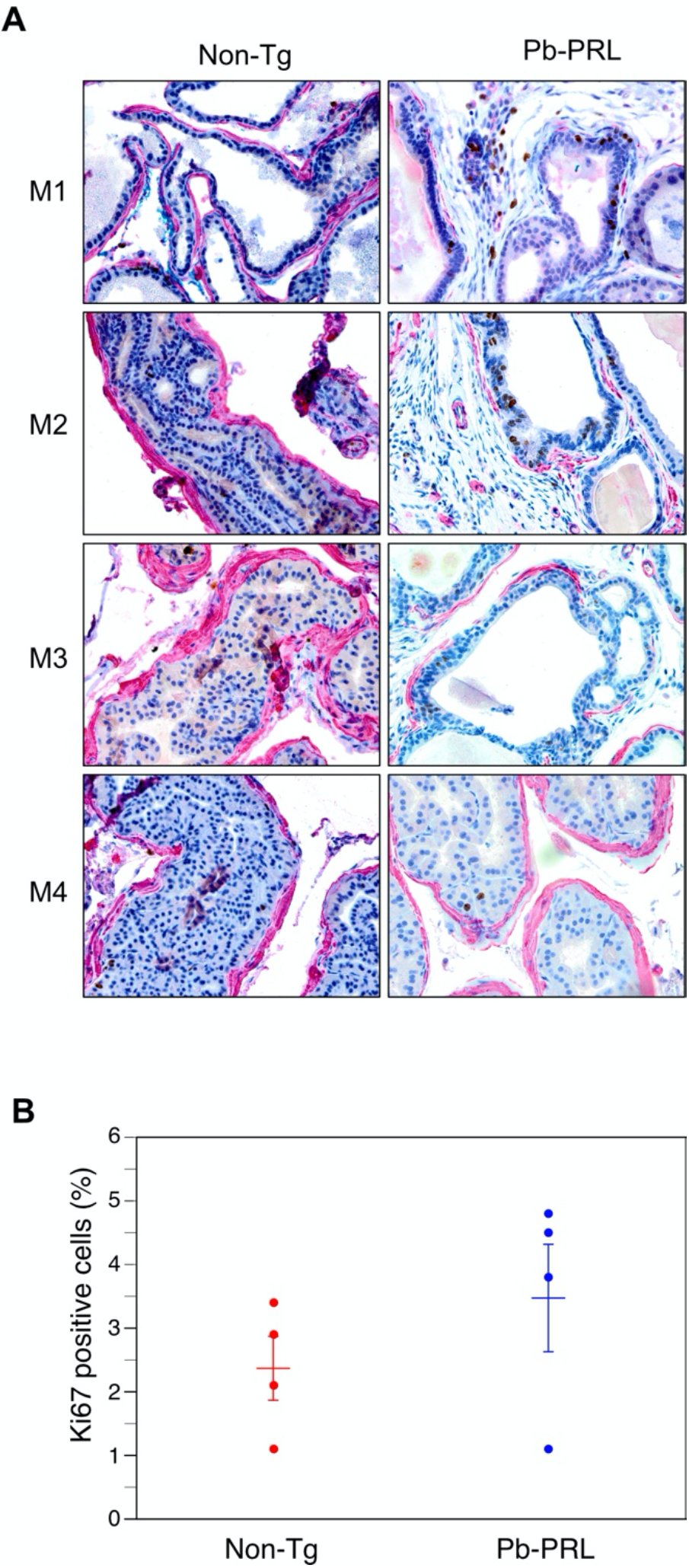
Prostate epithelium of Pb-PRL transgenic and non-transgenic mice. Non-transgenic (non-Tg) and Pb-PRL transgenic (Pb-PRL) mice were sacrificed and the prostates were fixed, sectioned, and IHC-stained with anti-Ki-67 and anti-αSMA, with hematoxylin as a counterstain. The epithelial glands are surrounded by α-SMA stained myofibroblasts. Ki-67-positive epithelial cells were counted in five high-power fields from each mouse prostate. **A.** Representative sections are shown for each mouse. Dual-staining for Ki-67 (brown) and α-SMA (red) in prostates from the indicated mouse strains. Magnification 400×. **B.** Quantification of Ki-67 positive cells in the epithelium of the prostates. Mean (+/-SEM) and individual values for each of four mice from each group is shown. Student’s unpaired t-test was used to compare non-Tg to Pb-PRL, p-value = 0.276.

We next examined the cellular composition of the stromal compartment. Specifically, we used anti-α-smooth muscle actin (anti-α-SMA) immunohistochemical staining and Masson’s Trichrome histochemical staining of collagen to visualize stromal components expressing the protein biomarkers representative of the two major cellular components of stroma: myofibroblasts and fibroblast, respectively. The anti-α-SMA immunostaining demonstrated that most of the myofibroblasts appear – on cross-section – to be in bands of stained-cells encircling the glandular structures in both the normal and BPH-like model mouse prostate (Fig. 5A; panels labeled ‘SMA’). There was no difference in the width or staining intensity of these bands, indicating no substantial differences in myofibroblast cell size and number between the Pb-PRL and the non-Tg littermate mouse prostates. The glandular epithelium of Pb-PRL mice were surrounded by dense stroma (Fig. 5A). In comparison, in the non-Tg littermate mice, the stromal compartment was relatively ‘sparse’ (with low nuclear density). Masson’s Trichrome stains both intra- and extra-cellular collagen blue, nuclei black, and myofibroblasts red. The intensity, cellularity, and the distance between epithelial-lined acini of the blue stained regions were all increased in the prostate tissue sections of Pb-PRL transgenic mice relative to those from the normal non-Tg littermate mice (Fig. 5A). This was confirmed by quantitation of the extent of collagen production, which is an indirect measure of the number of fibroblasts. The trichrome score, the product of the density and area of blue-colored tissue, was quantitated using imageJ as described [25], was increased in the Pb-PRL mouse prostate compared to the non-Tg prostate (Fig. 5B). Thus, the Pb-PRL BPH model shows an increase in the fibroblast-like component (including extracellularly-deposited collagen) of the stroma, but not in the myofibroblast component.

**Figure 5.**
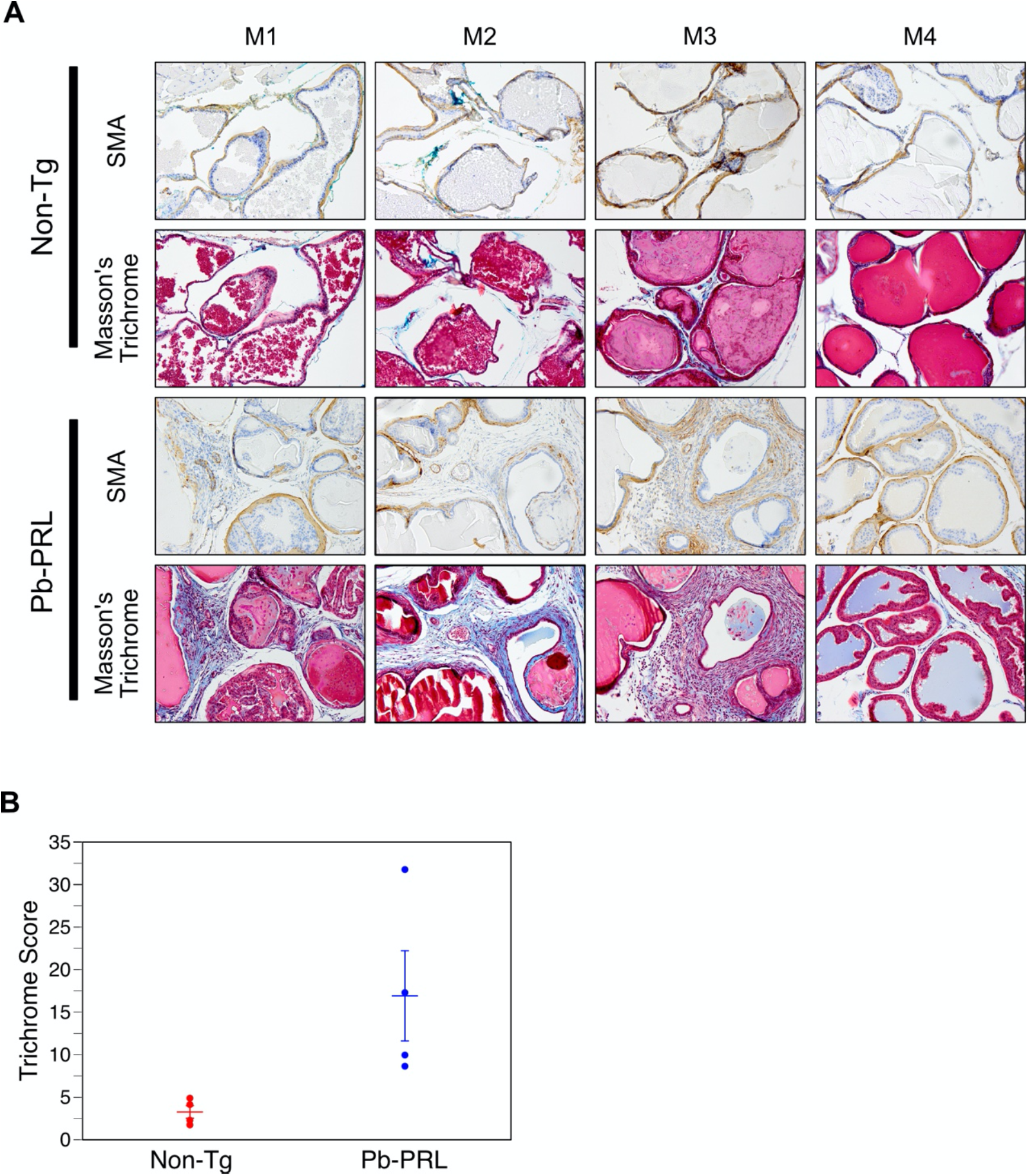
Increased stroma in prostates of Pb-PRL mice. Non-transgenic (Non-Tg) and Pb-PRL transgenic (Pb-PRL) mice were sacrificed and prostate tissue sections processed and stained. **A.** Images of immunostaining for α-SMA (smooth muscle actin, *brown*) and staining using Masson’s trichrome. Masson’s trichrome staining indicates collagen deposition (*blue*). Red staining indicates smooth muscle cells surrounding the glands (and non-specific red staining of the prostatic secretions intraluminally). Magnification 200×. **B.** Plots showing the mean (+/-SEM) trichome score in the stroma compartment for each of four mice of the two mouse strains. Note: Trichrome score is determined as described in Material and Methods section. Student’s unpaired t-test was used to compare non-Tg to Pb-PRL, p-value = 0.044.

### Regression of the stromal compartment in Pb-PRL prostates is reduced

Since Figure 5 indicates that fibroblasts and collagen deposition contribute to an increase in the size of the stromal compartment in the BPH model mouse prostate, we hypothesized that the rate of regression might be slower in the stromal compartment. Therefore, we measured regression rates for stromal and epithelial compartments, using manual 2D segmentation of the epithelial and stroma compartments in tissue sections, adjusted for residual prostate volumes using 3D HFUS reconstructions. Rates of regression for each compartment were then determined (Table 1). Castration induced regression in both epithelial and stromal compartment over time, for both the Pb-PRL BPH model and the non-Tg normal mice (Figs. 6 and 7, Table 1). In both non-Tg and Pb-PRL mice, the epithelial compartment (Figs. 6B, 7B) regressed faster than the stromal compartment (Figs. 6C, 7C). The rates of regression of the epithelial compartments were quite similar (Table 1). In contrast, the rate of regression of the stromal compartment in the Pb-PRL mice was more than 2-fold slower than in the non-Tg mice (Table 1). The difference in stromal regression rates between normal and Pb-PRL ventral prostate could account for the difference in prostate volume regression kinetics and residual volume observed in Figure 1. In both models, particularly the BPH model, the stromal compartment appeared more cellular 17 days after castration versus pre-castration, suggesting resistance to castration-induced cell death.

**Figure 6.**
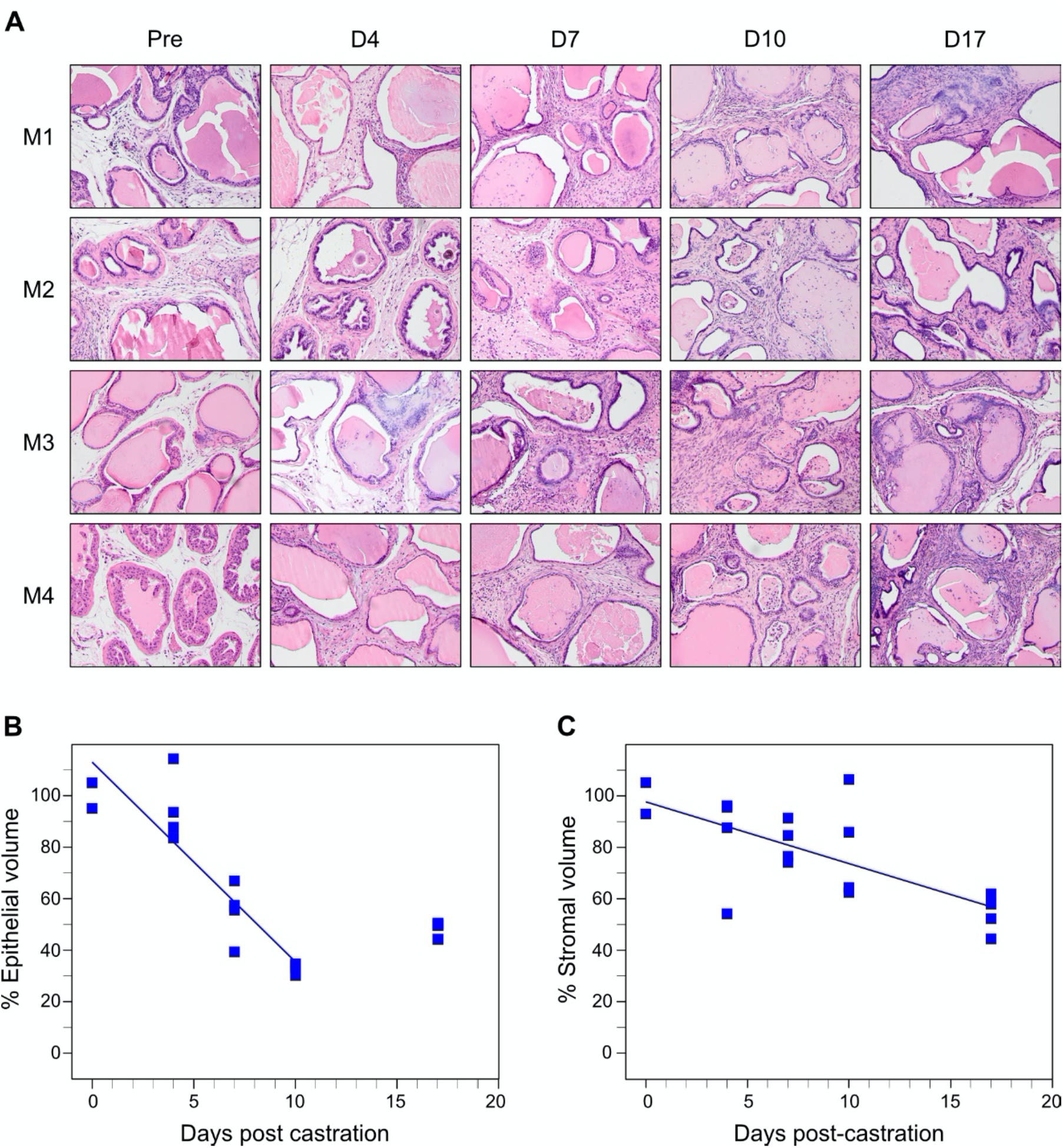
Castration induced regression in epithelial and stroma compartments in Pb-PRL transgenic mice. Mature Pb-PRL mice were surgically castrated and monitored by serial HFUS imaging to determine prostate VP volume for compartment area normalization (see methods). Prostates from four mice were examined at each of the five indicated time points following castration. **A.** Representative H&E stained sections correspond to the indicated times post-castration. Magnification 200×. **B.** Plot of normalized epithelial compartment volume percentage during ventral prostate regression. **C.** Plot of normalized stromal compartment volume percentage during ventral prostate regression. In **B** and **C**, each filled square represents an individual animal. Lines represent data fitted by least squares linear regression. Note that the epithelial compartment volumes determined at day 17 post-castration are not fitted.

**Figure 7.**
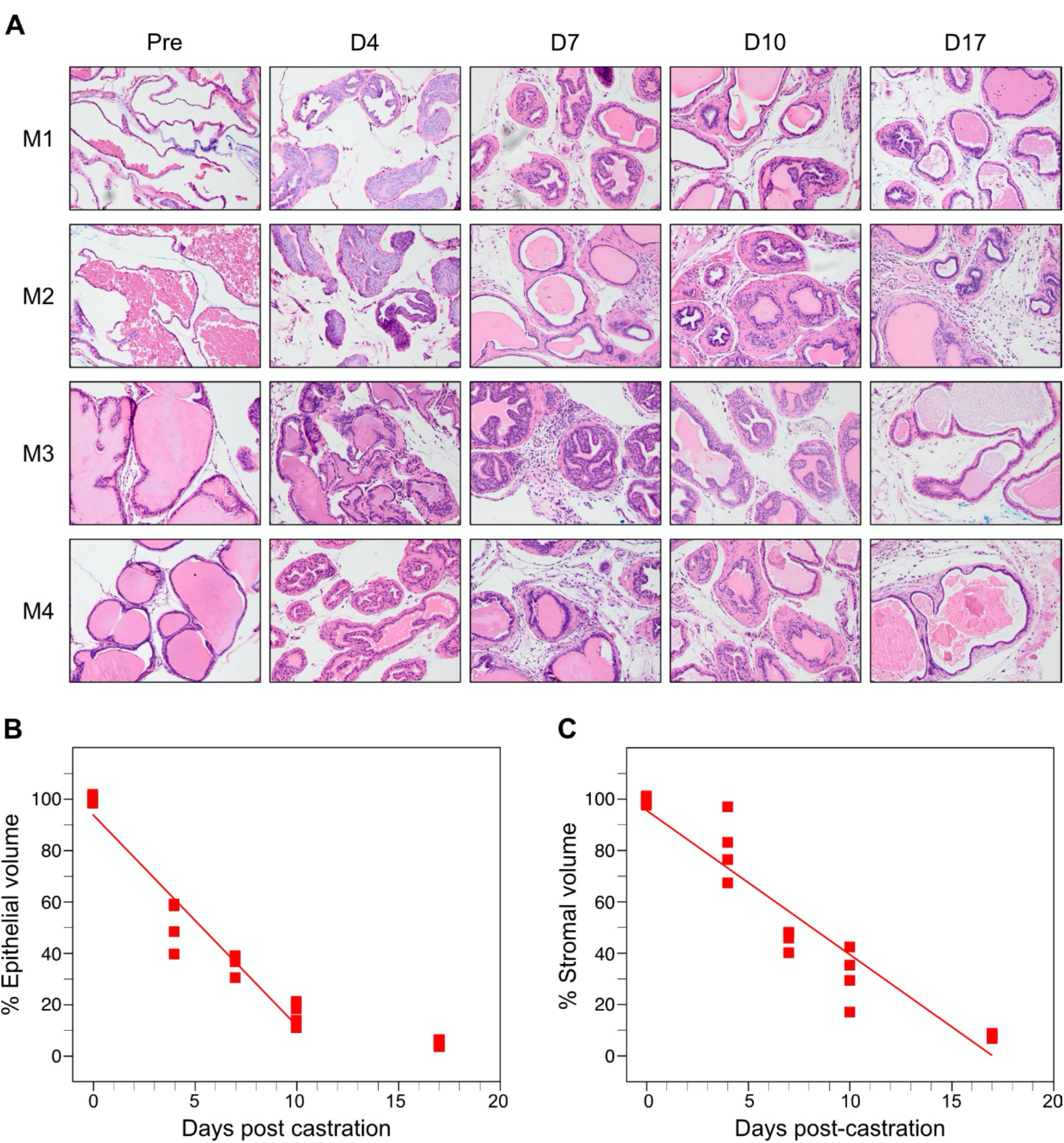
Castration induced regression in epithelial and stroma compartments in normal, non-transgenic mice. Mature littermate, non-transgenic mice were surgically castrated and monitored by serial HFUS imaging to determine prostate VP volume for compartment area normalization (see methods). Prostates from four mice were examined at each of the five indicated time points following castration. **A.** Representative H&E stained sections correspond to the indicated times post-castration. Magnification 200×. **B.** Plot of normalized epithelial compartment volume percentage during ventral prostate regression. **C.** Plot of normalized stromal compartment volume percentage during ventral prostate regression. In **B** and **C**, each filled square represents an individual animal. Lines represent data fitted by least squares linear regression. Note that the epithelial compartment volumes determined at day 17 post-castration are not fitted.

**Table 1.** Rate of castration-induced regression in epithelial and stroma compartments of normal C57Bl/6 or Pb-PRL mouse prostate. Note that the 17d time point was not included in the linear regression model for the two epithelial compartments, consistent with the plateau observed for epithelial regression in Figure 1. Regression slopes, 95% confidence intervals (CI) and R squared were calculated using a linear regression model. Stromal rate difference was tested using non-linear least squares regression modeling (F(2,34) = 34.0, p<0.001).

| Mouse/compartment | Rate of regression<br>(% of starting per day) | 95% CI | R squared |
| --- | --- | --- | --- |
| Pb-PRL Stroma | -2.401 | -3.696 to -1.107 | 0.4916 |
| Non-Tg Stroma | -5.615 | -6.510 to -4.720 | 0.9061 |
| Pb-PRL Epithelium | -7.761 | -10.09 to -5.465 | 0.8174 |
| Non-Tg Epithelium | -8.202 | -9.405 to -6.998 | 0.9385 |

### Cytokine signaling blockade following castration of Pb-PRL mice

Paracrine TNF and TGFβ1 signaling mediate castration-induced regression in both normal and neoplastic prostate [11, 17, 19, 20]. Therefore, we investigated the role of these multi-functional cytokines in hyperplastic prostates following castration, using the Pb-PRL model. Both TGFβ receptor phosphorylated Smads and the p38 mitogen-activated protein kinase (p38 MAPK) are important mediators of TGFβ-induced apoptosis [28]. To more completely inhibit TGFβ signaling, we administered both a TGFβR1 inhibitor and a p38 inhibitor prior to castration of Pb-PRL mice, and then twice a week thereafter, at a dose previously shown to block TGF signaling in mice [29, 30]. Similarly, we administered etanercept, a TNF signaling blocker active in mice [17] prior to and following castration. In both cases, HFUS was performed to monitor the volume of the VP. TGFβ-p38 signaling inhibition (red line) modestly retarded castration-induced regression in Pb-PRL mice over the first ten days following castration (Fig. 8A). However, TNF blockade showed little difference (yellow, Fig. 8B). Neither signaling inhibition of TGFβ nor TNF altered the overall extent of regression by 17 days after castration. These data suggest that paracrine signaling between the epithelium and stromal compartments by either cytokine was not a significant factor in the mechanisms that trigger regression in this BPH model.

**Figure 8.**
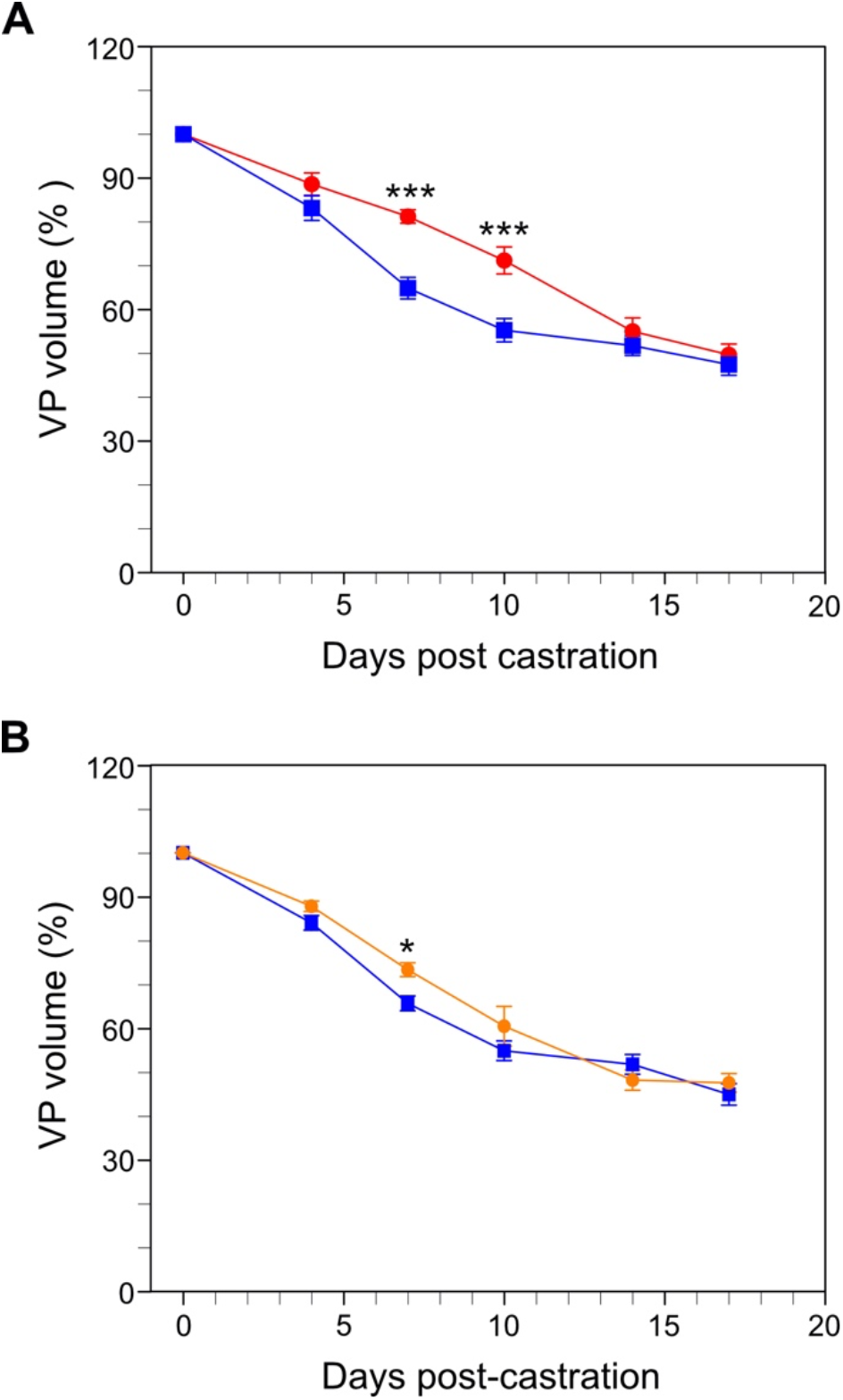
Blockade of cytokine signaling in Pb-PRL mice. Pb-PRL mice received vehicle (blue, data replicated from Fig. 1C), TGFβ/p38 signaling inhibitors (red) or TNF signaling blockade (orange) and were monitored after castration by serial HFUS imaging. **A**. Effect of TGF-β (red) signaling blockade on castration-induced regression of VP. **B.** Effect of TNF (orange) signaling blockade on castration-induced regression of VP. Data points represent mean volume relative to starting volume, +/-SEM (N=4). \**P* <.05; \*\*\**P* <.001. TGF-β or TNF-α inhibitor treated castrated mice were compared to Pb-PRL tg castrated mice using two-way ANOVA with Tukey’s HSD test.

## Discussion

In these studies, we employed Pb-PRL mice, arguably the best murine model of BPH to better understand androgen withdrawal signaling of prostate regression. Prolactin (PRL) and its cognate receptor (prolactin receptor; PRLR) are expressed in both human [31] and rat prostate [32]. In the first attempt to produce a prolactin-driven model of BPH, transgenic PRL was expressed under the control of a metallothionein-1 (Mt-1) regulated promoter were first generated to study the role of prolactin (PRL) in the prostate gland [33]. Prostates became enlarged but did not progress to cancer. However, PRL increased the levels of circulating testosterone, complicating interpretation of this model [34]. The androgen-regulated rat probasin (Pb) promoter has been widely used to produce targeted prostate epithelium-specific transgene expression. In a subsequent refinement of the model, Pb-PRL transgenic mice engineered to overexpress the rat PRL gene under the control of the Pb promoter developed a dramatic enlargement of the prostate gland [22, 35]. Notably, no high-grade PIN or tumor formation was detected in the Pb-PRL prostate, consistent with well-documented concept that human BPH is a hyperplastic and hypertrophic disease, that bears no direct or even indirect linkage to prostate cancer. Unlike Mt-PRL mice, Pb– PRL mice do not exhibit elevated serum androgen [22]; thus, Pb-PRL mice are likely to be a better model of human BPH development and of BPH response to therapy. Unfortunately, there is currently no evidence that PRL plays a significant role in the initiation or maintenance of BPH in humans [36]. However, consistent with the hyperplastic phenotype of human BPH, we observed a remarkably uniform, sigmoidal growth pattern for the Pb-PRL prostates that plateaued at about 10- or 20-fold the volume of the adult normal VP. In contrast, the PTEN-loss mouse model of prostate cancer exhibited a variable, exponential prostate (tumor) growth pattern (refs. [20, 37] and Figure 1).

BPH is androgen dependent, and in the Pb-PRL model prostate weight is halved when stromal AR is knocked out [21]. Our HFUS imaging enables accurate serial monitoring and real-time analyses of prostate volume kinetics in live mice both developmentally and following castration. As shown in Figure 2, we observed a sizable residual volume following castration in the Pb-PRL BPH model relative to normal prostate in non-Tg mice. Morphometric measurements in Figures 6 and 7 (summarized in Table 1) demonstrate that a reduced rate of regression of the stromal compartment – relative to the epithelial compartment – accounts for the residual volume. This essentially represents a failure of the stromal compartment to fully regress. Figure 5 suggests that the residual volume consisted largely of increased collagen (both intracellular in fibroblasts as well as extracellular deposition) in stromal compartment of the Pb-PRL prostate. Given the almost 10-fold increase in the Pb-PRL VP volume relative to the normal VP, it is likely that a substantial portion of the residual volume is due to an expanded population of fibroblasts and concomitant extracellular collagen. Notably, the data in Figure 5 also indicates that there was little expansion of the relative number of myofibroblasts, except perhaps surrounding the increased number of glandular elements. One limitation of these studies is the limited duration of volume and histological assessment after castration, and future studies might explore extended treatments.

Gustafsson and colleagues performed a detailed morphological assessment of human BPH [38]. Sections from 16 BPH patients were stained with steroid receptors (ERα, ERβ1, ERβcx, and AR), proliferation markers (Ki-67 and PNCA), the antiapoptotic factor Bcl-2, cytokeratin 8 (CK8), and EMT/ TGFβ-mediated transcription factors (pSmad3, Snail, and Slug). These investigators do not detect a significant amount of stromal proliferation in human BPH, which is consistent with the very low Ki67 staining pattern we observed in the stromal compartment in Figure 4. Instead, they propose that an epithelial-mesenchymal transition (EMT) contributes to the stromal accumulation in human BPH. Specifically, they suggest the trans-differentiation of epithelial and endothelial cells into stromal fibroblasts. Each of the TGFβ ligands (β1, β2, and β-3) are mediators of epithelial and endothelial to mesenchyme transition [39, 40]. Cell supernatant from normal prostate stromal WPMY-1 cells was able to induce EMT in BPH-1 cells by secreting TGF-β1, which activated Smad3 signaling [41]. In addition to EMT, TGFβs are also critical drivers of proliferation inhibition and apoptosis induction [42], including prostate cell apoptosis [27, 43, 44]. When we administrated inhibitors of two TGFβ signaling mediators, TGFβR1 and p38 MAPK, Pb-PRL VP volume decreased at a modestly slower rate for the first 10 days following castration (Figure 8A). The reduced sensitivity of trans-differentiated fibroblasts to castration-induced TGFβ-driven apoptosis may account for the reduced rate of regression of the stromal compartment of the BPH model.

Our previous studies have demonstrated that TNF is necessary for castration-induced prostate regression and that castration acutely increases TNF levels in the prostate [27]. In the normal mouse, the prostates from Tnf^-/-^ and Tnfr1^-/-^ mice regressed considerably more slowly after surgical castration, and the regression in Tnf^-/-^ prostate could be restored by administering soluble TNF [17]. Etanercept blockade of TNF signaling recapitulated the effects of genomic loss of Tnf. While TNF was required for castration-induced regression of normal prostate, neither FasL nor TRAIL were necessary [17]. In contrast to the normal prostate where TNF blockade reduces regression within four days [17], TNF signaling blockade did not affect the rate or extent of castration-induced regression of the Pb-PRL mouse prostate over 17 days (Fig. 8). We reported prostate cancer cells rely on TRAIL to mediate castration [45], and this suggests that other death receptor ligands may be active inducers of castration-induced apoptosis in the Pb-PRL model of BPH.

Nonetheless TNF plays an important role in BPH. The mechanism of TNF-mediated promotion of prostatic hyperplasia is unknown, but inflammation-driven proliferation is likely [10]. We recently reported long-term (12 week) TNF signaling blockade reduced ventral prostate volume in Pb-PRL mice by diminishing epithelial cell proliferation [46]. Anti-TNF therapies reduced macrophages and NF-kB activity in both BPH patients and in non-obese diabetic (NOD) mice (which have autoimmune inflammation-associated prostatic hyperplasia). This suggests that TNF is a viable therapeutic target in men with BPH. Indeed, the incidence of BPH is reduced in men treated with anti-TNF drugs [46].

Our data indicated that finasteride is less effective than castration (Fig. 3) thus suggesting that therapies that reduce androgen levels might improve the treatment of BPH. Luteinizing-releasing hormone (LHRH) antagonists, which reduce prostate volume by lowering testosterone to castrate levels, showed promise in men with BPH/LUTS in a Phase II, randomized, placebo-controlled study, but Phase III trials involving over 1,600 patients failed to reduce their International Prostate Symptom Score (IPSS) [8]. The trials were terminated even though a third Phase III safety study Z-041 with 528 patients reported a nearly 6-point reduction in IPSS. Nonetheless, LUTS is effectively relieved by androgen deprivation therapy with LHRH antagonists in men with prostate cancer [18, 47], suggesting LUTS in this population is multi-factorial. The residual stroma in hyperplastic prostate after castration indicates standard ADT is an unlikely future therapy for men without neoplastic epithelia driven LUTS. Since finasteride both dramatically reduces prostate volume in this BPH model (Fig. 3), and has been shown to reduce both prostate volume and recurrence of LUTS in men with BPH [48], early long term 5ARIs in combination with alpha-adrenergic receptor blockers may be cost effective [49] and the most efficacious therapy.

## Acknowledgements

This work was supported by grants HHS-6-15SF and HHS-009-17SF from the S.A.S. Foundation, 126771-IRG-14-194-11 from the ACS, the Roswell Park Alliance Foundation (all KLN); the Department of Defense Prostate Cancer Research Program No. W81XWH-19-1-0378 (JJK), and the National Cancer Institute Cancer Center Support Grant P30-CA016056 to Roswell Park Comprehensive Cancer Center.

